# Low-Rank Tensor Encoding Models Decompose Natural Speech Comprehension Processes

**DOI:** 10.1101/2025.06.02.657514

**Authors:** Lane Lewis, Xaq Pitkow, Leila Wehbe

**Author notes:** denotes equal contribution on senior authorship (Xaq Pitkow, Leila Wehbe).

## Abstract

How does the brain process language over time? Research suggests that natural human language is processed hierarchically across brain regions over time. However, attempts to characterize this computation have thus far been limited to tightly controlled experimental settings that capture only a coarse picture of the brain dynamics underlying human natural language comprehension. The recent emergence of LLM encoding models promises a new avenue to discover and characterize rich semantic information in the brain, yet interpretable methods for linking information in LLMs to language processing over time are limited. In this work, we develop a low-rank tensor regression method to decompose LLM encoding models into interpretable components of semantics, time, and brain region activation, and apply the method to a Magnetoencephalography (MEG) dataset in which subjects listened to narrative stories. With only a few components, we show improved performance compared to a standard ridge regression encoding model, suggesting the low-rank models provide a good inductive bias for language encoding. In addition, our method discovers a diverse spectrum of interpretable response components that are sensitive to a rich set of low-level and semantic language features, showing that our method is able to separate distinct language processing features in neural signals. After controlling for low-level audio and sentence features, we demonstrate better capture of semantic features. Through use of low-rank tensor encoding models we are able to decompose neural responses to language features, showing improved encoding performance and interpretable processing components, suggesting our method as a useful tool for uncovering language processes in naturalistic settings.

## 1 Introduction

Natural language processing happens over time and regions in the brain, from early auditory processing to word parsing to semantics. Characterizing how language is processed has largely been carried out using non-invasive recording modalities namely, fMRI, EEG and MEG. Of these methods, MEG boasts a better temporal resolution than fMRI and better spatial resolution than EEG, making it an ideal candidate to study the fast processing of language. Classical literature has found stereotyped responses in brain activity to language in highly controlled experiments, such as the N400 response [1]. However, these experiments typically only focus on one aspect of language processing at a time and might not reflect how the system behaves in naturalistic settings.

As an alternative to the highly controlled setting, language experiments with naturalistic stimuli have become increasingly common in MEG [2, 3, 4]. To analyze these complex data, researchers have paired encoding models with word and sequence embeddings, often derived from language models. Recent research has identified a rich relationship between the processing of large language models (LLMs) and natural language processing in the human brain. Advances in LLMs have proven to be a source of rich language features that can successfully predict human brain responses [2, 5, 6, 7, 3]. Conversely, the human brain has also served as inspiration for language and speech models, demonstrating an ability to drive their improvement in some cases [6, 8, 9] [10]. However, due to the lack of interpretability of the complex embeddings of language models, it is not simple to make sense of these correspondences. Further, because the stimuli are not controlled by design, different low-level features can be correlated with semantic features represented in the embeddings, leading to spurious language model performance. Thus, it is difficult to determine which language features in an LLM lead to good predictions of brain activity. Recent research has suggested that language model features can be confounded with low-level features and simple model architecture changes such as position encoding [11]. This highlights the need for new encoding model methods to be developed that can better disentangle the features of language model components contributing to brain activity, to better separate low level confounds from meaningful language processing.

In this paper, we use a natural audio stimulus, and propose a principled way of extracting the relationship between language model features over time and sensors using a low-rank Canonical Polyadic (CP) tensor encoding models We use this method with a magnetoencephalogrophy (MEG) dataset where subjects listen to naturalistic narrative stories to propose a more accurate encoding model that relies on a low-rank constraint with fewer parameters to train more generalizable models than full-rank MEG encoding models. These low rank models also serve to to better understand speech comprehension processing. We demonstrate that our model is able to separate out the contributions of low-level auditory features, such as ends of sentences, from higher-level semantic features such as pronouns or emotion, and these distinct contributions have some spatial and temporal separation.

## 2 Related Work

### Encoding Models

Encoding models are defined as a map from a of feature set of stimuli to brain activity (often restricted to be linear) with the goal of predicting neural activity [12, 13][vision paper, language paper]. These models have been used in a variety of contexts and modalities, such as vision [14, 15, 16, 17, 18, 19], audition [20], and language [21, 22, 23]. For modeling language responses, a popular and performant choice of features is to use a pretrained neural network language model [2, 5, 6, 7, 3], which host a battery of rich features useful for predicting a range of brain measurements including MEG [2, 6, 3]. These learned embeddings enable a new paradigm of neuroscience, allowing models to effectively describe neural activity during naturalistic behaviors and generalize outside of trained stimuli, motivating their use as digital twins [24, 13]. However, interpreting encoding models is fraught with difficulty. One salient challenge is how to separate low-level and high-level processing features, which limits their use in providing neuroscientifically useful results [11]. Our work provides a new method for building encoding models that naturally separates low-level and high level language processing using low rank relationships over time and sensors.

### Encoding Model Interpretation Methods

Understanding which stimuli features contribute to brain prediction performance in complex encoding models can be difficult. One method is variance partitioning, which can quantify the unique variance predicted by different feature groups [25, 26, 27, 28], but it relies on knowing the groups of features in advance. One common method of characterizing an encoding model is to find the most activating stimuli for a particular brain region. This method has shown a degree of generalization, correctly predicting stimuli outside of the original stimulus set that when presented back to the brain strongly drive neural responses [29, 30, 31, 32]. An alternative method is to examine the weights on which features best predict neural activity. In fMRI, this approach has successfully identified semantic areas of the brain after projecting encoding model weights on their top principal components [23, 19] or by using matrix decomposition on the brain data [33]. Comparatively less interpretation work has been done with encoding models for MEG [3, 34, 4], which can deliver important knowledge about the dynamics of language processes

### Low-Rank Regression Models

Tensor regression models are a natural extension to linear regression when independent and/or dependent variables have a multilinear structure. Similarly there exist tensor extensions of low-rank regression to different notions of low rank tensors, such as restricting the Tucker Tensor decomposition or Canonical Polydiadic (CP) Tensor decomposition [35, 36]. Neural activity has been found to lie on low dimensional subspaces in a variety of experimental settings and across brain areas [37, 38]. This has motivated the use of low rank methods to find structure in neural activity, including tensor decompositions and tensor regressions [39, 40]. Low-rank tensor regressions have also been used in neuroscience for regression problems [41, 42]. While low-rank *matrix* encoding models have been used in language neuroscience [9] to facilitate training (without interpretation), low-rank *tensor* methods have so far not been explored.

## 3 Methods

### 3.1 Low-Rank Tensor Encoding Model

We define a time-delay encoding model on MEG as the following: Let the data of each story *i* be represented as the MEG signal 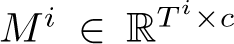 collected over *T* ^*i*^ time points and *c* channels, with an additional tuple set 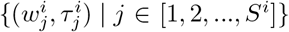 denoting the story words, 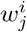, spoken at corresponding time, 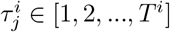 with a total number of words *S*^*i*^ occurring in each story. For each word, the LLMs provide an embedding 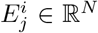. We construct our embedding time-series dataset as zero everywhere except at word times 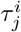:

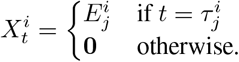

Unlike fMRI, whose time course is slower than the usual pace of spoken words, MEG is fast enough to resolve brain activity during the processing of each word. To transform from word embeddings to our MEG signal, we use a linear 3D FIR filter model with *D* time delays and *C* MEG sensors. This filter *F* ∈ ℝ^*D*×*N* ×*C*^ is convolved with the time series defined by the embeddings and their timings, and predicts activity at every sensor. This gives us a linear encoding model for predicting MEG responses 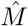 where

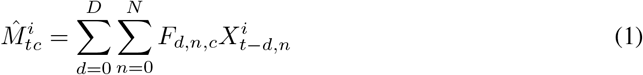

To build a low rank tensor encoding model, we construct *F* as a low rank Canonical Polydiadic (CP) tensor. We define a tensor filter of rank *R* as the sum of *R* rank-1 filter components, each given by a product of three factors selected as rows of the matrices *U*^*D*^ ∈ ℝ ^*R*×*D*^, *U*^*E*^ ∈ ℝ ^*R*×*N*^, and *U*^*C*^ ∈ ℝ ^*R*×*C*^ denoting time delay factors, embedding factors, and channel factors (see Figure 1):

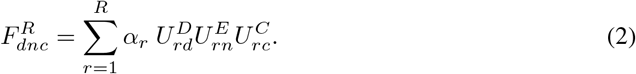

**Figure 1:**
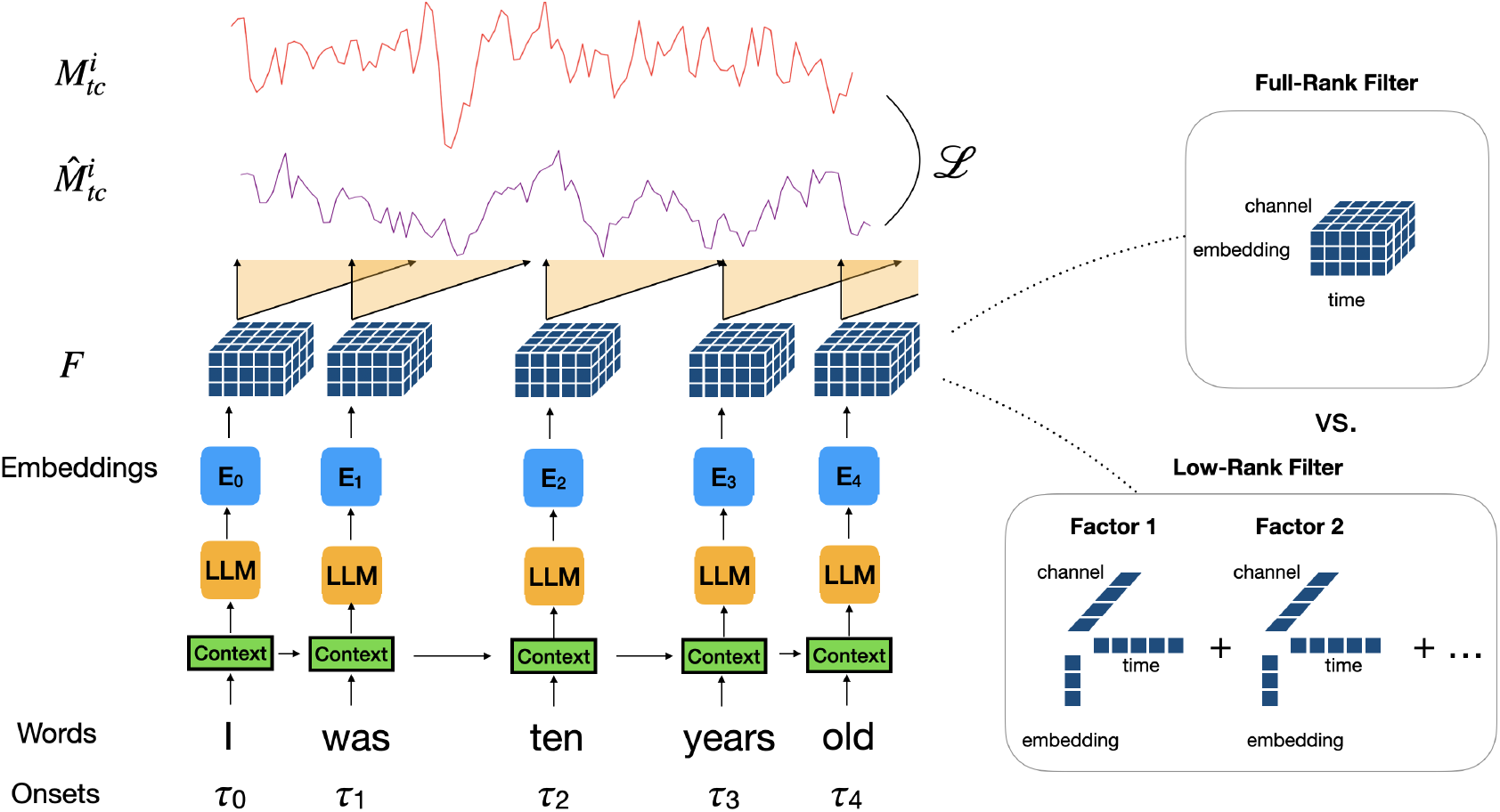
Low-rank tensor encoding model predicts brain activity along distinct rank-1 filter weight components over time, channels and language model embeddings. The previous 20 words provide context for the LLM embeddings, and the word onsets give the timing of the embedding, giving a high-dimensional time series. The low rank weight filters are then convolved with this time series and summed over components to generate the prediction of each channel over time.

Each factor is normalized to length 1; all scaling is absorbed into one coefficient *α*_*r*_ per component *r*.

### 3.2 Model fitting

We define our training loss on the MEG training data to be the mean squared error (MSE), with an optional ridge regularization parameter *λ*_*c*_ for each channel to account for per-channel differences in signal-to-noise:

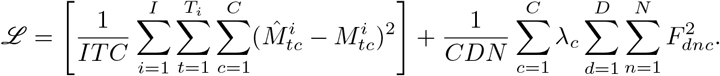

where 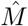 is given by Equation 1.

To fit the full rank tensor model, we use ridge regression where the ridge penalty parameters were chosen per channel using 6-fold cross-validation on an exponential parameter set from [0.001, 10^4^]. To take into account our model’s correlation structure over time, we split the dataset into folds composed of sequential MEG data of equal time lengths within each story. This ensured that our *K*-folds’ training and validation data remained distinct. During fitting, we do not restrict the factors to be norm 1. Instead, post-fitting, the normalization constant is pulled out from the factors.

To fit the low-rank tensor model given computational constraints, we trained to convergence using stochastic gradient descent with the Adam optimizer, a batch size of 300,000, random initialization, and a learning rate of 5 × 10^−3^. Given that each channel is no longer fit independently like in full linear regression, correct regularization requires a combinatorial search over ridge hyperparameters per channel, but this is computationally infeasible. Instead, we choose the ridge penalty to be 0.1 for all channels, near the average channel hyperparameter of a full linear regression model. To build the models we use the pytorch library [43] and fit each low-rank run using a mix of 20 CPUs with 512GB RAM and an L40 or A6000 GPU on a local HPC cluster. Each low-rank tensor model takes around 3 hours to fit, with an estimated 200 GPU hours of used to generate the results in this paper and 80 extra hours used for early exploratory runs and analysis.

In order to control how language features flow through our models, we needed to choose a context window size and the number of time delays to use. Similar to previous research, we choose a standard context window size of 20 words [34]. We choose 40 time delays so that we can capture activity up to 800ms after word onset on our 50Hz sampled MEG data, where we expect most of the language-driven signal to lie [34].

To generate the language model embeddings, we use Llama-2 7b. Previous research has found that layer 3 is the best layer for predicting MEG activity during natural speech [44], so we choose that layer for all subsequent analysis. To reduce our computational burden, we take advantage of previous work showing LLM embeddings lie in low-dimensional subspaces [45]. We therefore run PCA on the 4096 dimensional embeddings over our training set, and restrict our analysis to the 665 dimensions that capture 95% of the variance (Supplemental Figure 1), which we use for the rest of our analysis.

To measure the performance on our held-out repeated test story we use the *CC*_norm_, which calculates the correlation of the test data’s repeat mean response over the estimated noise ceiling. To remove poor estimations of the noise ceiling on low-signal channels, we set the minimum noise ceiling as 0.2. The noise ceiling was calculated using the method from Schoppe, Oliver, et al.[46, 47]

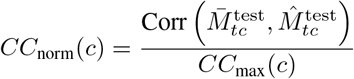

where 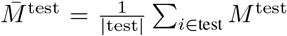 is the average response over the five repeats of one story, and *CC*_max_(*c*) is the estimated noise ceiling on the test correlation for a channel.

### 3.3 MEG Dataset and Preprocessing

To understand the processing of natural language over time, we chose to use an MEG dataset with natural language stimuli but also natural timing. In this experimental setup, subjects listened to naturalistic stories in the form of podcasts from the Moth Radio Hour. These stories have already been studied in fMRI [48], and have been later scanned in MEG by an anonymous research group. In total, datasets for 3 subjects were collected using 27 unique stories over 5 sessions. Within each session, one story was repeated twice. Additionally, one story was repeated across all sessions. The data was collected in a MEGIN scanner using 306 channels with 102 cranial points at 1khz. Participants gave their informed consent and were remunerated for their time, and the study was approved by an anonymous institutional review board.

For MEG scan preprocessing, external sources were removed from the MEG signal with SSS filtering, and a notch filter removed 60Hz signal from the power lines. The data was then band pass filtered between 1hz and 150hz. Finally, ICA was run to remove any sources highly correlated to eyeblinks or the raw audio signal. During initial analysis, we noticed that the audio was stretched by a small (imperceptible) factor while being played in the scanner. This stretch was approximately consistent within each story across subjects; we corrected for this effect by resampling. All analysis was performed on a version of the MEG signal that we downsampled to 50hz for computational efficiency. In total our dataset comprised of approximately 58,000 words (excluding the words from repeated stories) and approximately 1.3 million MEG samples per subject.

The labels of words and their respective timings were used from [48]. However, the words consisted of capitalized and un-punctuated words which are likely to be out of distribution for a language model. To build a transcript that falls into natural language, we used a strong language model, GPT-4, and prompted it to keep the transcript words the same but convert it into our desired format. After passing it through the language model, each transcript was human checked and repaired to ensure that each word remained the same.

For all training runs, we used the same dataset composed of all stories except for a held out test set of one new story with 5 repeats. This ensured no contamination between training and test. Before training, the MEG data was normalized per channel per story.

## 4 Results

### 4.1 Neural Predictability Post Word Onset

Before fitting encoding models to our dataset, we wanted to investigate the predictability of the neural signal across channels. To do this, we used neural responses to 5 repetitions of one story to compute the noise ceiling, *CC*_max_. Subjects displayed predictability in a variety of brain regions (Figure 2c, Supplemental Figure 2), with more predictability in auditory and language processing regions.

**Figure 2:**
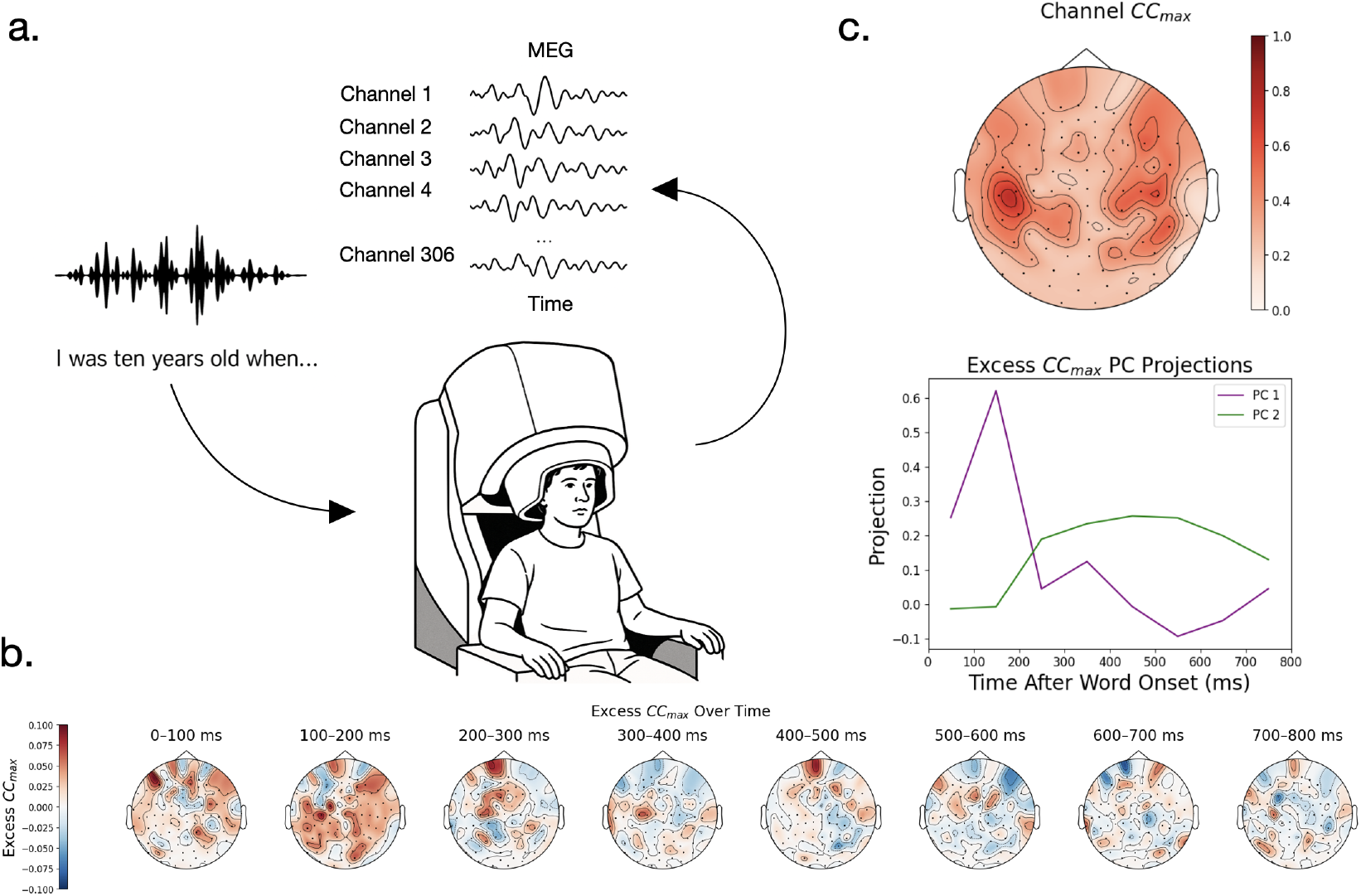
a) Subjects listened to Moth Radio Hour stories in an MEG scanner. The rest of the plots are shown for subject A. b) The predictability after word onset minus the average predictability (excess *CC*_max_) estimates over channels show distinct timecourses over, responding strongly 100-200ms and again more weakly at 400-500ms indicating there are predictable time courses in the neural data post-word onset. c) Top panel: Neural activity shows higher predictability around the auditory cortex and language processing regions. Bottom panel: Projecting the excess *CC*_max_ onto its first two Principal Components (PCs) over channels reveals that after the auditory onset response, neural activity should be predictable on the timescale of semantic processing.

To see how the predictability of the signal changes as a function of time after word onset, we made subsets of our neural data over time and computed their noise ceiling. Each subset was constructed using a time window post-word onset of 100ms. These time windows were then concatenated together to form a subsampled time series which we used to compute the *CC*_max_. To view the change in the difference of *CC*_max_ over time, we subtract out the *CC*_max_ over all time points to get the excess *CC*_max_ given by that time window.

We find that the excess *CC*_max_ fluctuates over time and channels (Figure 2b for subject A, other in Supplemental Figure 3), with a distinct peak in 100-200ms for all subjects, likely corresponding to initial processing of the audio signal, and a shallower, broader peak again around 400ms in subjects A and D, likely corresponding to more semantic processing. Doing PCA on the excess *CC*_max_ of subject A makes these timescales more apparent. These analyses suggest that predictability of neural activity varies on multiple timescales post-word onset (Figure 2a).

### 4.2 Low-Rank Encoding Models Improve Performance

A low-rank encoding model carries an inductive bias that the data being modeled is low dimensional. This might be relevant to MEG data either because the underlying brain activity is low dimensional or because, due to physical limitations and distortions, the MEG sensors sample from a broad region of underlying cortex, thus leading to redundancies. To investigate whether the low-rank inductive bias is useful in predicting MEG activity during natural language processing, we fit a series of low-rank models with increasing rank (Figure 3a). We find that indeed low-rank models predict neural activity better, in some cases drastically such as Subject A, where the encoding performance *CC*_*max*_ effectively doubles. To test the significance of this result, we perform a bootstrap hypothesis test against the performance of the full regression model as well as a permutation test against chance performance. We find all performance to be significant at 0.001 except for Subjects C and D’s rank 1 models, which fail to significantly outperform regression (Supplemental 11, 12). To see where low-rank models drive encoding performance, we took rank-10 models and compared their performance over channels to the full-rank model (Figure 3c, Supplemental Figure 4). We find that the difference is generally widespread, improving the predictions over many areas. These results suggest that low-rank models are more predictive than standard encoding models on our MEG dataset.

**Figure 3:**
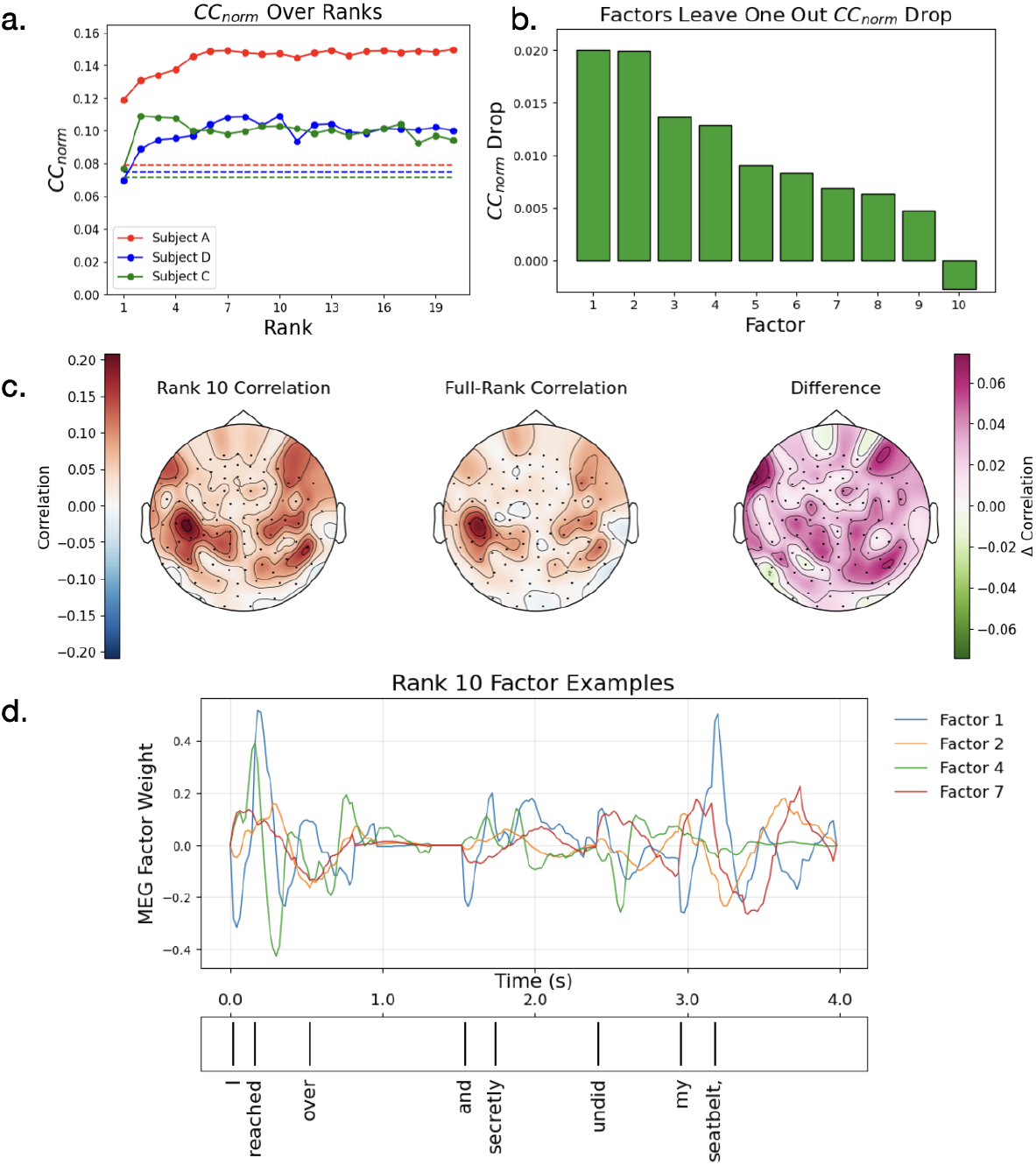
a) Low-rank tensor models improve encoding performance, measured by the channel average *CC*_norm_. Dotted line represents the performance of a full rank regression model for that subject. Low rank performance saturates early with a small number of components. b) Components contribute differently to encoding performance, measured by the drop in performance when omitting that component. c) Low-rank models improve encoding performance against full models broadly over channels (subject A). d) Timecourses of component weights: Each component is driven by a pattern of language features over time, and predicts a spatial pattern across channels according to 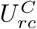 (Equation 1). These component weights (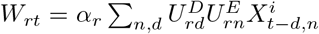) indicate the strength of each of these spatial patterns in the predicted MEG signal. These components of the brain response to language show a diversity of processing over factors (subject A).

### 4.3 Low-Rank Components Capture a Diverse Spectrum of Neural Activity

We now ask what language processes the low-rank model is capturing. We find that low-rank encoding models learn diverse, complex activations over time in response to language features (Figure 3d, Supplemental Figure 5). However, their contributions may not be equally predictive of the MEG signal. To estimate the predictive power of a single factor, we consider the drop in mean *CC*_norm_ when that factor is excluded from prediction. We find that not all factors contribute equally, with some factors having a much larger effect than others and, in rare cases, dropping factors leading to improvements (Figure 3b, Supplemental Figure 6). For subsequent analysis, we sort components by their leave-one-out influence, starting with the most influential component.

Next, we study the factors individually. Since our factors are defined as having a vector norm of 1, with the norm value being absorbed into a single multiplicative coefficient *α*_*r*_, we have no scale ambiguity in our interpretations (Equation 2). We find that the coefficients are roughly the same across components even though they have different influence (Supplemental Figure 11, Figure 3b). To analyze the components, we examine a model with 10 components for all subjects. Within this low-rank model are a diverse set of processing time-courses and spatial modes (Figure 4, Supplemental Figure 7). Largely we find that the spatial modes sit over auditory and language areas (Figure 4, Supplemental Figure 8). The timecourses vary but many show peaks around 150–200ms and 300–400ms.

**Figure 4:**
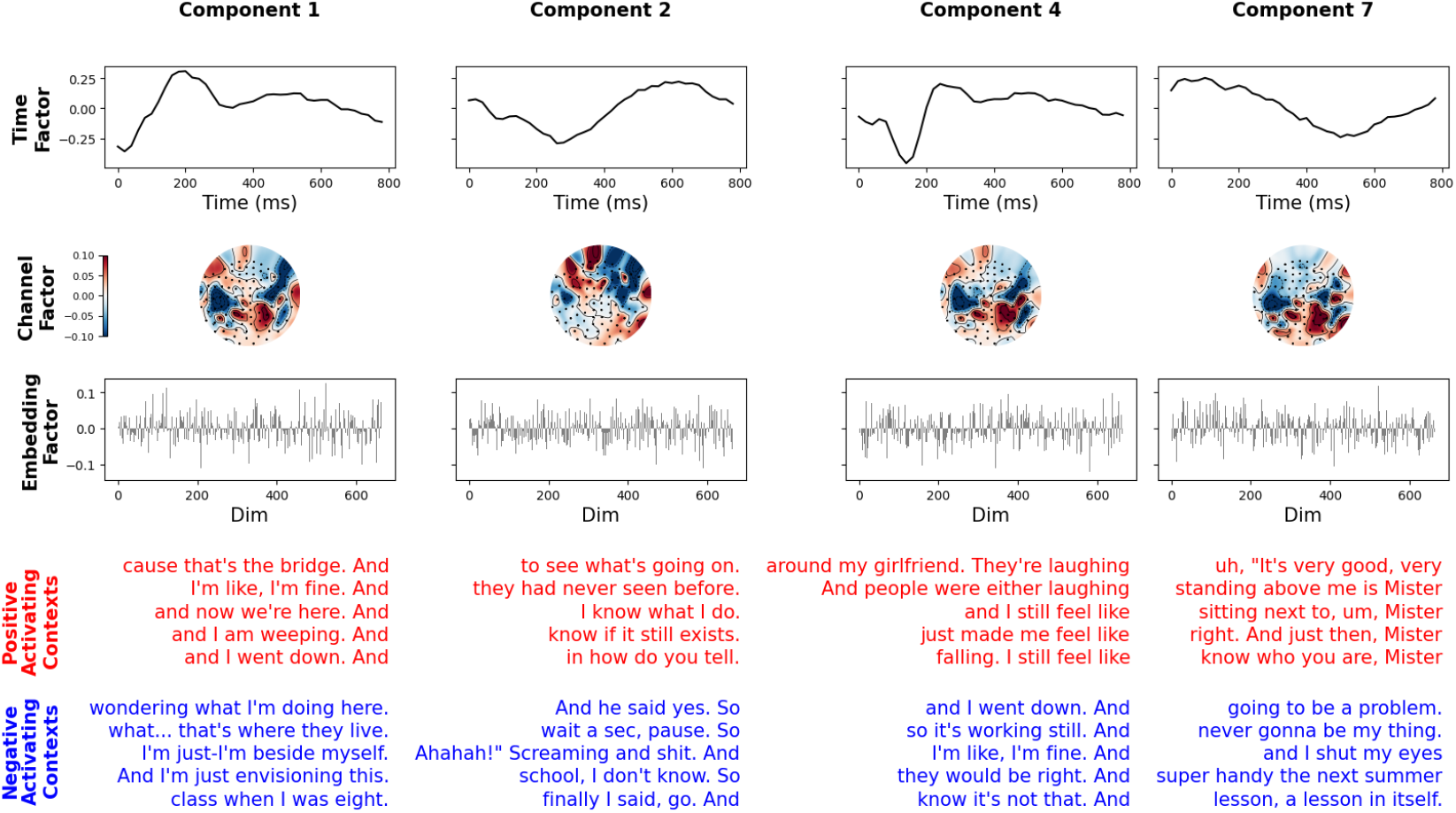
The low rank model decomposes neural activity into distinct 1-dimensional factors that depend on time, channels, and language embedding. *Top three rows*: Several example components’ factors are shown from a rank-10 model of subject A, demonstrating the spectrum of different activations over time and channels. Several time factors and channel factors show distinct patterns, suggesting the decomposition is separating distinct aspects of language processing. *Bottom two rows*: Top 5 most positively (red) and negatively (blue) activating five-word contexts, for each component’s language embedding factor. The factors are largely dominated by low level effects such as sentence start and end (components 1,2,4) while some capture more semantic information (component 7).

To understand the language features driving each component, we searched over the space of contexts from the Moth Radio Hour transcript corpus. MEG predictability saturates after just a few words of context, consistent with previous literature [34] (Supplemental Figure 9). Because of this, we examine the most activating contexts with 5 words. For each component we select the five most activating contexts from all transcribed stories as those whose LLM embedding have the highest dot products with that component’s embedding factor 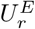. Similarly, for the most negative activating contexts we select the most negative dot products. Examining these selected contexts, we find that most factors are activated by low level language features, such as end of sentence and start of sentence (Figure 4, factor 1). However, some respond to more semantic features (Figure 4 factor 7). To quantify the low rank features, we count the ratio of factors whose most activating (positive or negative) components respond to only starts of sentences or punctuation. We find that 8/10,9/10 and 9/10 factors of subjects A, C and D satisfy this criteria.

In summary, the low-rank components capture a wide set of distinct language processes over time, channels, and language embeddings. These components are dominated by the encoding of low-level features but high level aspects also contribute to MEG prediction. In the next section, we control for the effect low-level features and show that the model can still predict high-level components in MEG.

Anecdotally, our model was quite useful at detecting an experimental artifact. In development, we noticed that some of our factors were nonsensical and showed high-frequency temporal fluctuations. This allowed us to diagnose some leakage of audio-signal into the MEG signal, which we later fixed in preprocessing. This example showcases the utility and power of our low-rank decomposition.

### 4.4 Low-Level Controls Improve Semantic Features

We previously found that the low-rank components are dominated by low-level effects. We want to see if subtracting out these low-level signals leads to any changes in the semantic content captured by the low rank factors. For our control set, we constructed a features space of a log-mel spectrogram, word onset and sentence start/end. To construct the log-mel spectrogram, we downsample each story audio signal to 16kHz with a hop length of 25ms and 80 Mel features over 15 time delays. We encode word onsets, sentence starts, and sentence ends each as binary sequences taking a value of 1 at the times the respective features appear, and 0 at all other times. We then fit a ridge-less full-rank model over the train and test set of the features and regressed out the control signals.

After controlling for these low-level language properties, we notice several changes in the components of the residual activity. First, they shift towards more semantic features in their most activating contexts (Figure 5, Supplemental Figure 10) with only 3/10 still encoding low-level features in subject A and 8/10 in C. Subject D showed less of this trend as it maintained 9/10 low level language features. Second, through using a normalized power analysis over factors, we show the peaks of the spatial factors shift away from the original auditory regions toward other brain regions (Figure 5b, Supplemental Figure 10,13). Third, using the same power analysis, we show the timecourses shift, peaking at later times, consistent with longer processing times for higher level language comprehension (Figure 5a, Supplemental Figure 14).

**Figure 5:**
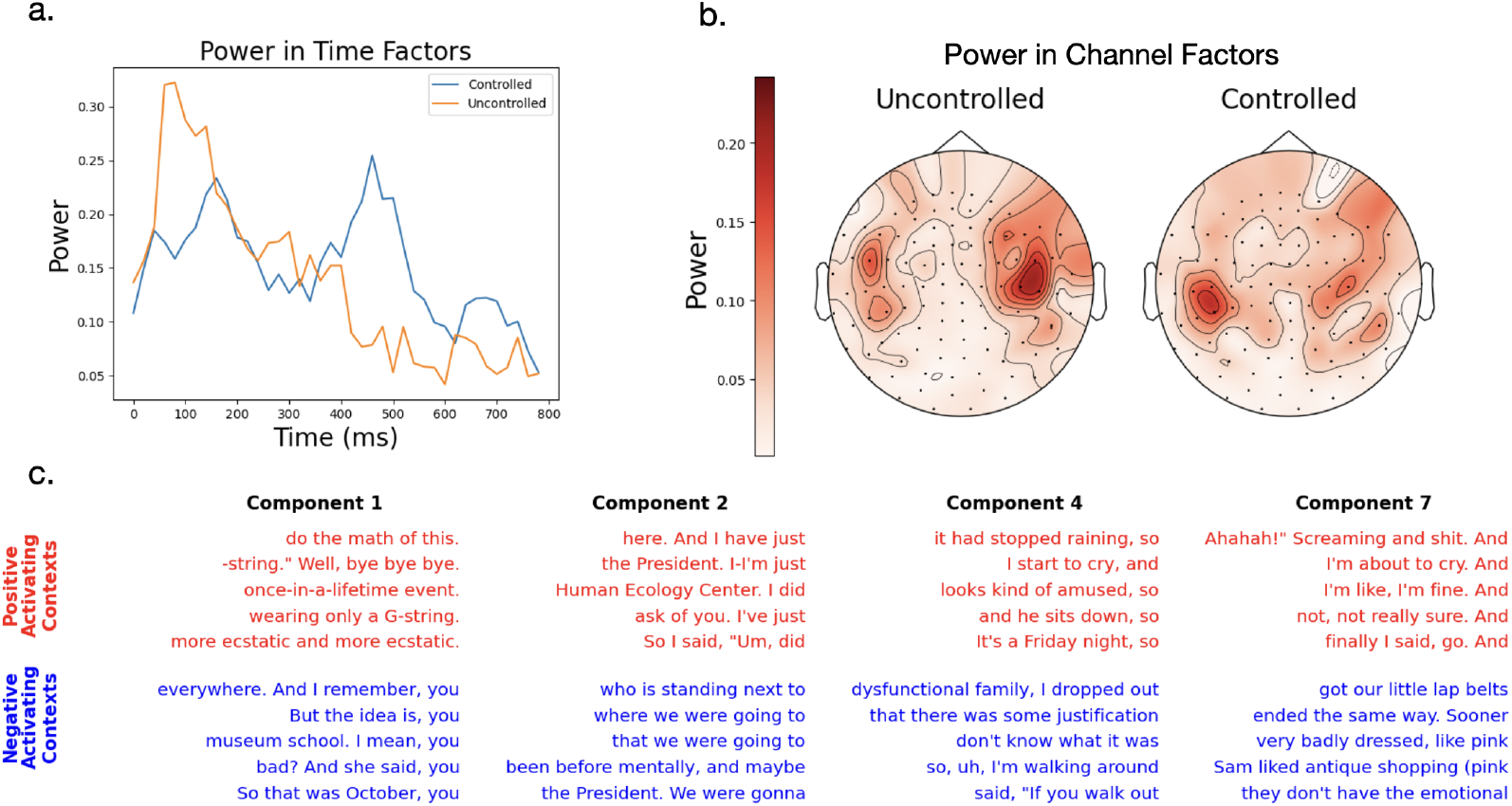
Low-rank model of residual responses after controlling for low-level language features, compared to model applied to original uncontrolled neural activity. All panels shown for Subject A. a) : Power in time factors shifts later in to residual responses versus uncontrolled data. Un-normalized power here is defined as 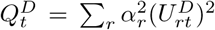 and the power is *P*^*D*^ = *Q*^*D*^*/*||*Q*^*D*^||.b) Power in channel factors becomes more distributed for residual data versus uncontrolled data. Un-normalized power here is given by 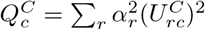 and power is *P*^*C*^ = *Q*^*C*^*/*||*Q*^*C*^||. c) Words that activate the factors’ language embedding most (red) and least (blue), as in Figure 4. These contexts capture more semantic and less low-level features like ends of sentences.

Thus, after removing low level controls, we find more semantic features are captured in the low rank components. This indicates that though the presence of low-level confounds is a concern for understanding language model brain predictability, low-level features do not capture all model predicted variance. Our results suggest that future encoding model works should focus on capturing explainable variance after subtracting out low level controls to capture meaningful language processing.

## 5 Conclusion

In this work we predicted brain activity from language features using low-rank tensor encoding models. We showed that this method improves encoding performance over full rank linear regression models. Beyond this, we demonstrate that our model is able to naturally and effectively capture interpretable low- and high-level language processing features over time and space. Finally, by controlling for low-level features, we show we can isolate more semantic components.

While we applied this low-rank tensor encoding model to an MEG dataset, we see it as a general method useful for building interpretable and performant encoding models in neuroscience. We are excited about the application of this method to other encoding model modalities like video and audio as well as other brain recording methods such as fMRI, EEG, and microelectrode recordings.

While our model is able to capture diverse language processing features in multiple subjects, our analysis was limited to a comparative study over each subject individually. To better capture the low-rank components shared across subjects, a single low-rank model could be used on all subjects, stacking MEG features over subjects listening to the same auditory stimuli. Additionally, in this paper, we subtract a limited number of controls from the MEG signal when investigating the model’s learned semantic feature encoding. To improve the interpretations of the model factors, more rich control sets could be used. We leave both these as lines of investigation for future work.

## 6 Funding and Competing Interests

X.P. and L.L. were supported in part by the National Science Foundation and DoD OUSD (R & E) under Cooperative Agreement PHY-2229929 for the NSF AI Institute for Artificial and Natural Intelligence, ARNI. This work was also partially supported by the National Institute On Deafness And Other Communication Disorders of the National Institutes of Health under Award Number R01DC020088. The content is solely the responsibility of the authors and does not necessarily represent the official views of the National Institutes of Health. The authors declare no competing interests.

## Notes

### Competing Interest Statement

The authors have declared no competing interest.

