## Supplemental Figures for "Low-Rank Tensor Encoding Models Decompose Natural Speech Comprehension Processes"

### Supplemental Material

May 22, 2025

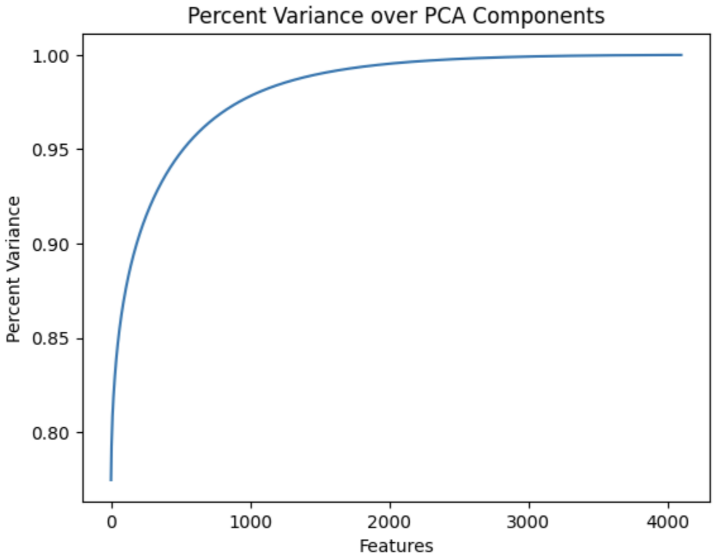


Figure 1: Embedding percent variance explained over PCA components of Llama2-7b on the Moth Radio Corpus. 95% of the variance is captured at a small percent of the total dimension.

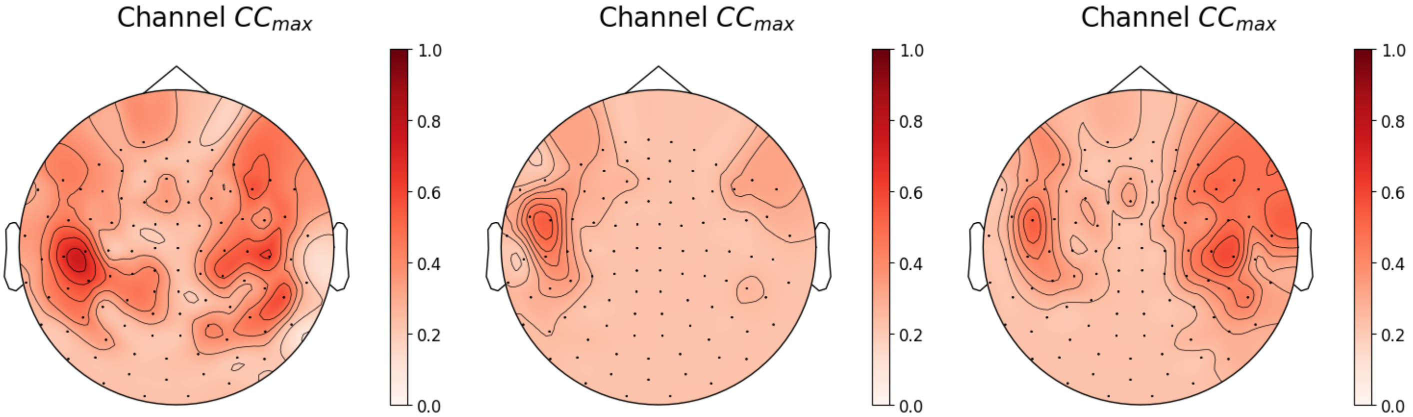


Figure 2: $CC_{max}$ over channels. Subjects A and D show much better predictability over channels. *Left*: Subject A. *Middle*: Subject C. *Right*: Subject D.

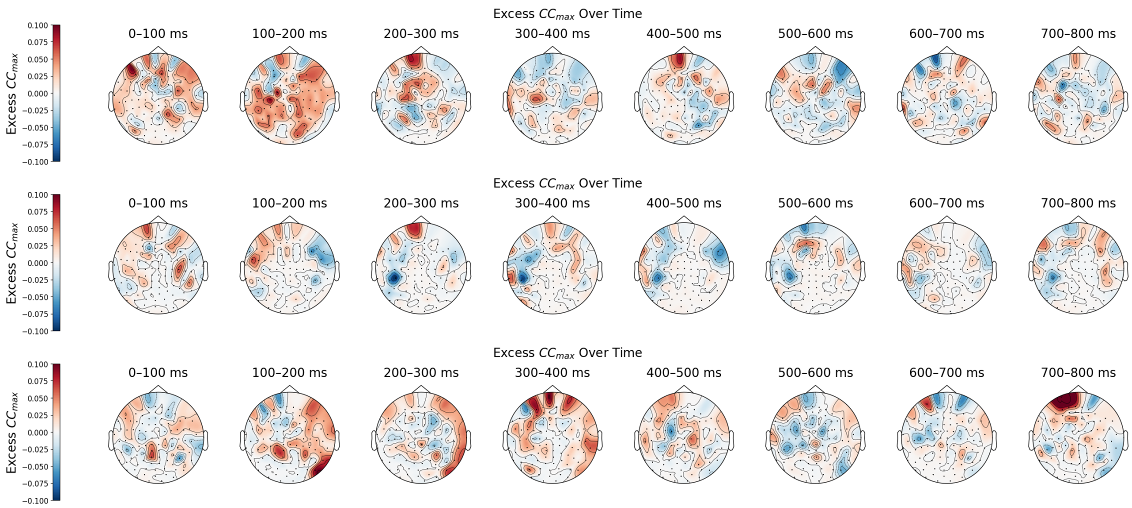


Figure 3: Channel excess $CC_{max}$ shows distinct patterns over time. Subjects A and D show early peaks in predictability around 100-200ms post word onset and an additional shallow peak at 400-500ms in subject A and 300-400ms in subject D. *Upper panel*: Subject A. *Middle panel*: Subject C. *Lower panel*: Subject D.

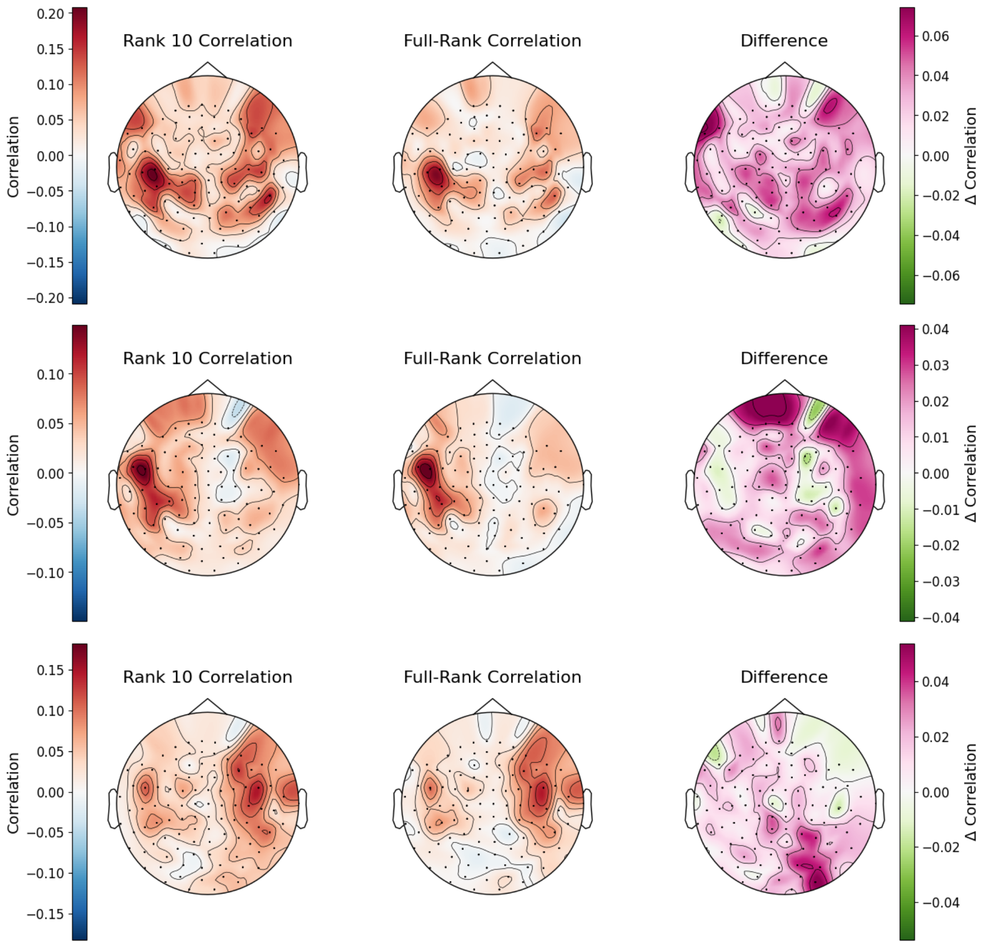


Figure 4: Rank 10 models show widespread performance improvements over full ridge regression. *Top panel*: Subject A. *Middle panel*: Subject C. *Bottom panel*: Subject D.

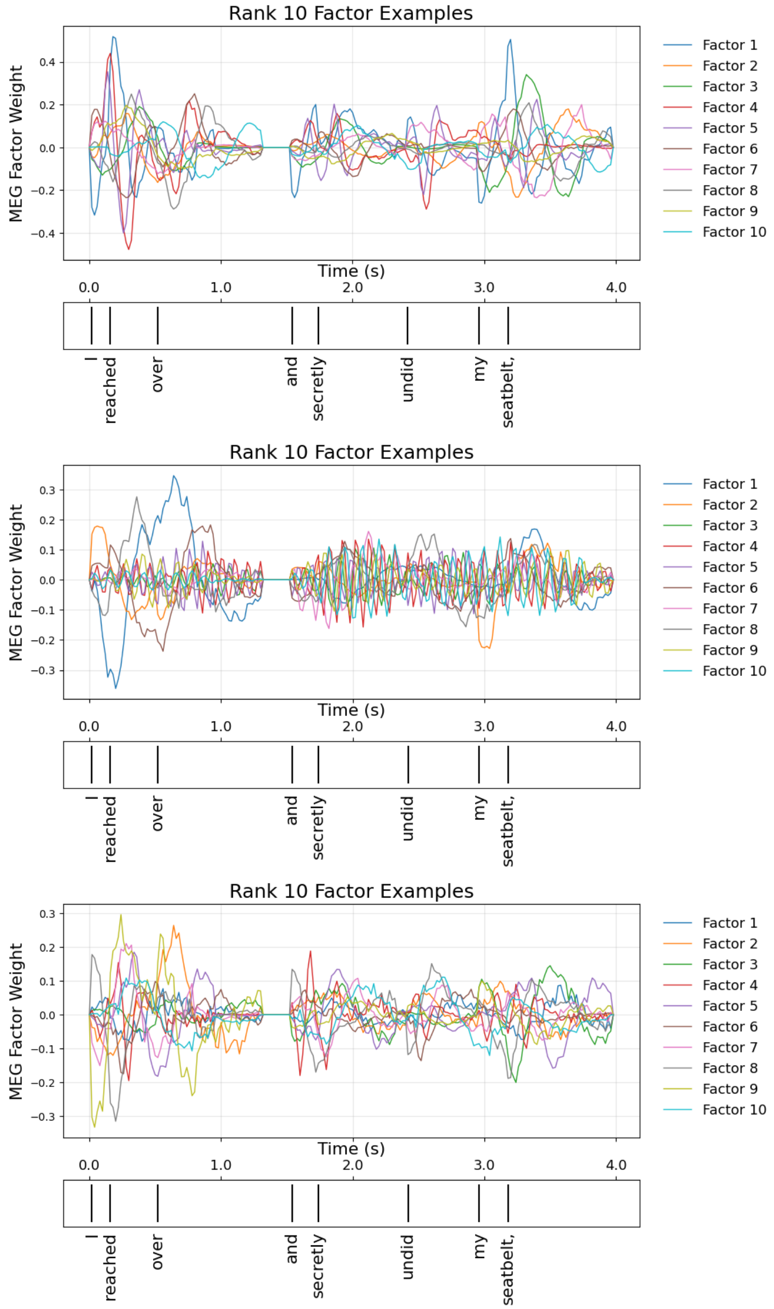


Figure 5: Timecourses of components pre-MEG factor multiplication. Factors show a diversity of timecourses in response to natural language. *Top panel*: Subject A. *Middle panel*: Subject C. Note the very fast components, likely corresponding to the small amount of signal in subject C compared to the number of ranks. *Bottom panel*: Subject D.

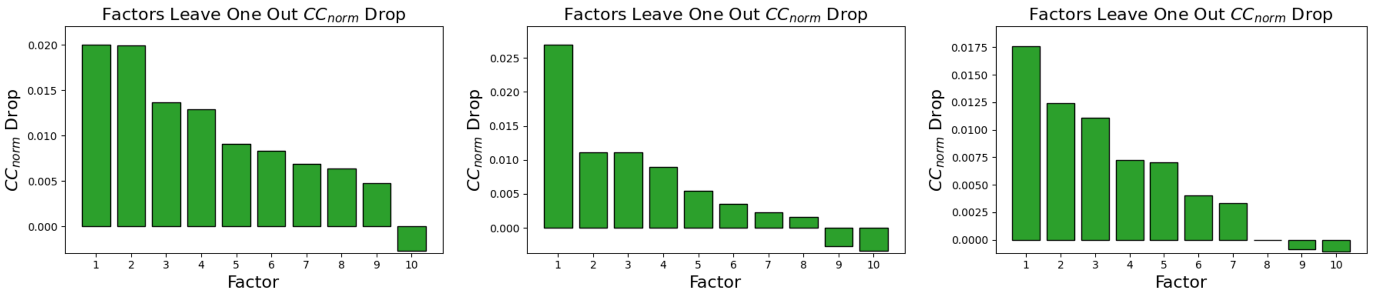


Figure 6: Leave one component out drop in $CC_{norm}$ over rank. Components show different influence on test performance. *Left*: Subject A. *Middle*: Subject C. *Right*: Subject D.

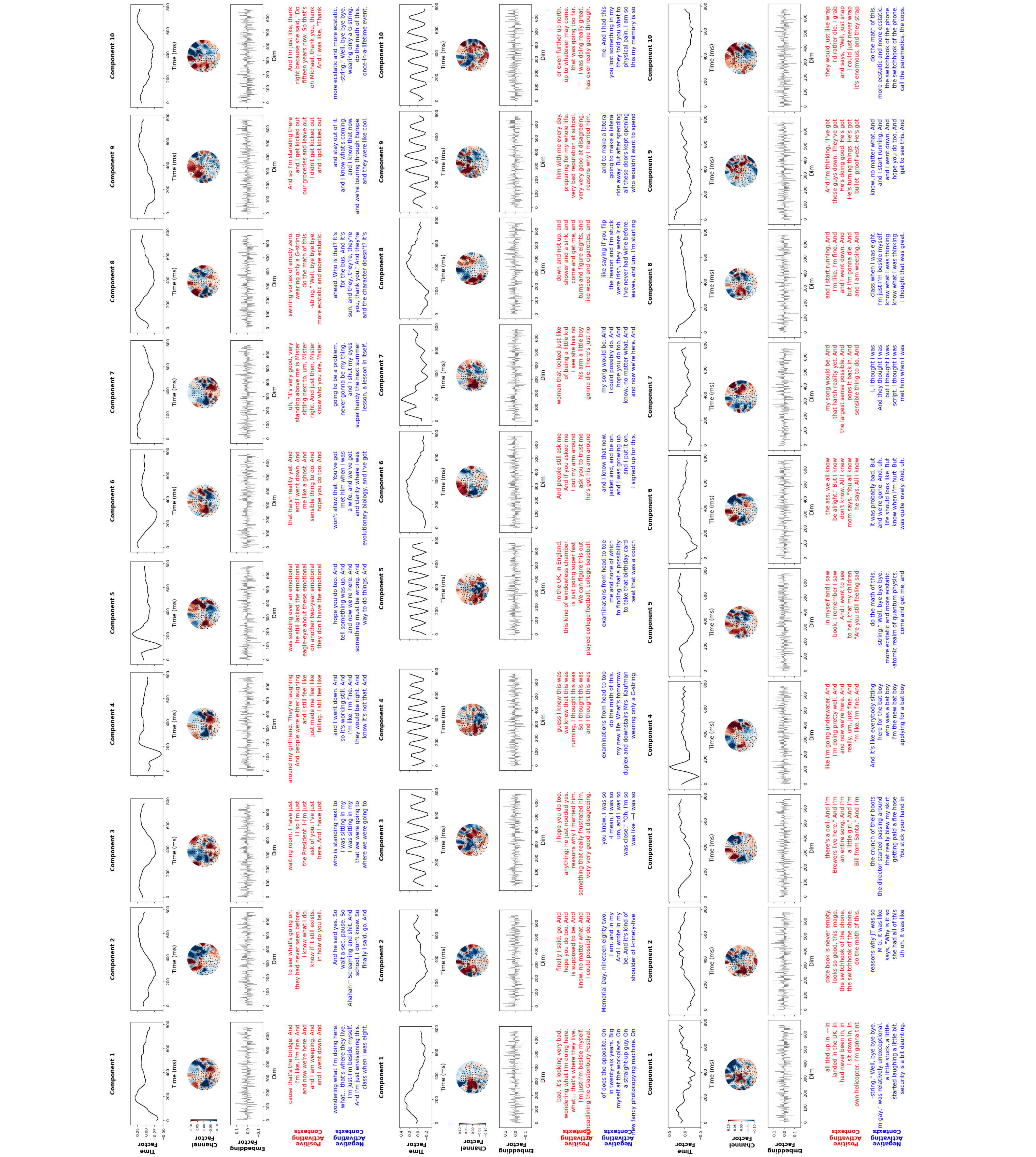

Figure 7: Subjects show a diversity of timecourses, spatial components, and activating texts. *Left panel*: Subject A. *Middle panel*: Subject C. Note the fast timecourse components, likely stemming from low signal in the subject. *Right panel*: Subject D.

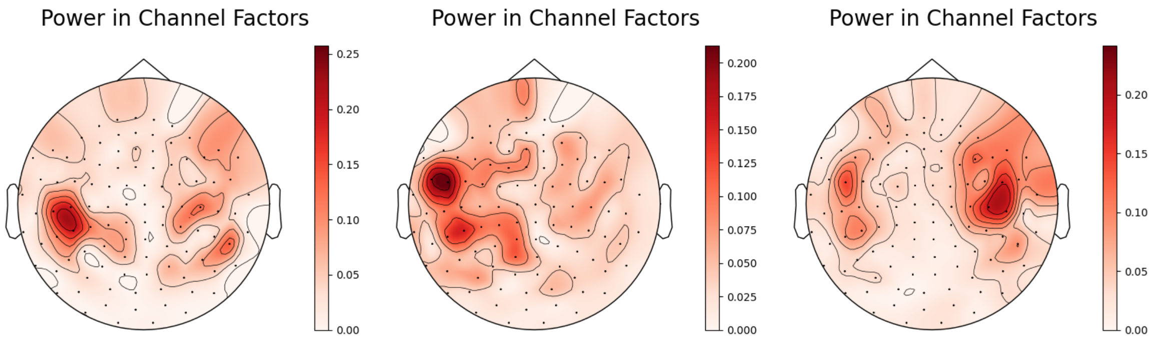


Figure 8: Using the same power analysis as main figure 5 shows that subject channel factors concentrate over auditory areas (left and right hemisphere, roughly in temporal and prefrontal regions).*Left*: Subject A. *Middle*: Subject C. *Right*: Subject D.

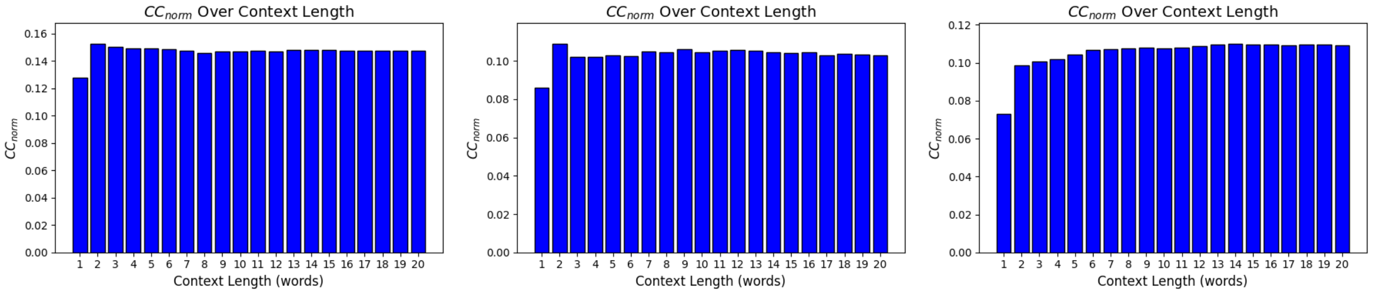


Figure 9: $CC_{norm}$ saturates after a small number of previous words being included in the context when evaluated on a rank-10 model trained on a 20 word context window. *Left*: Subject A. *Middle*: Subject C. *Right*: Subject D.

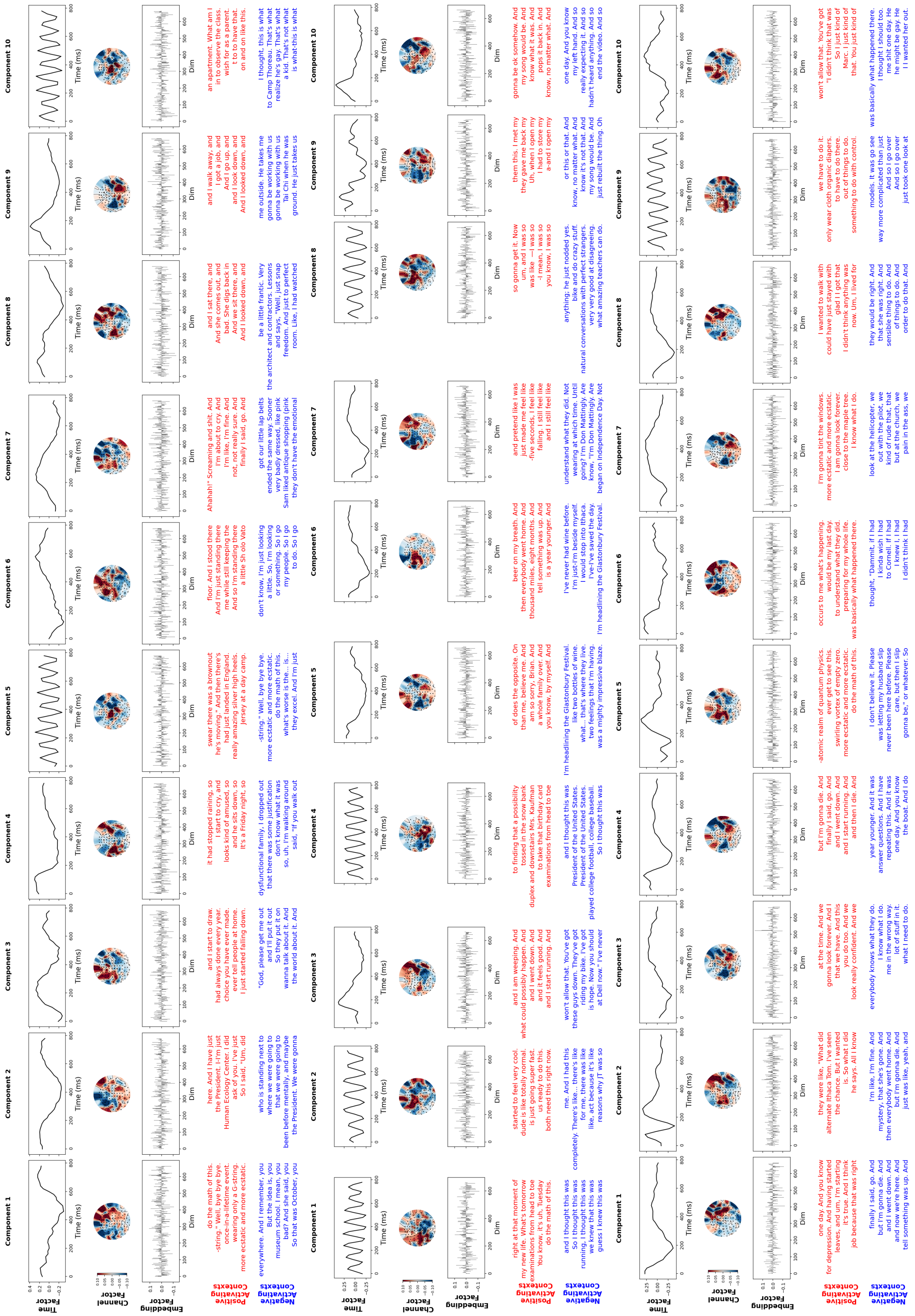

Figure 10: Control subtracted factors for a rank-10 model. We see changes in timecourses, channels and activating texts from the un-controlled rank-10 model. Note the fast timecourses, likely arising from less signal in control subtracted residuals. *Left panel*: Subject A. *Middle panel*: Subject C. *Right panel*: Subject D.

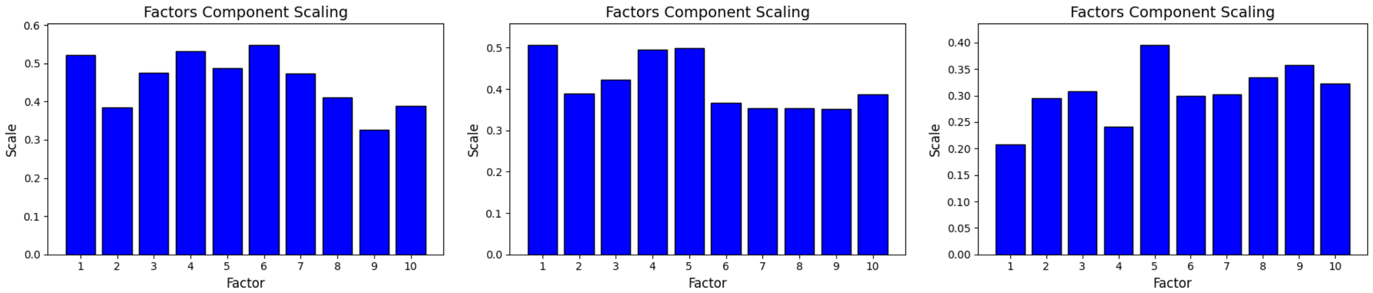


(a) Component scalings show roughly similar ranges over components despite having different influences on test performance (see Supplemental Figure 6).*Left*: Subject A. *Middle*: Subject C. *Right*: Subject D.

| Subject | Rank 1 | Rank 2 | Rank 3 | Rank 4 | Rank 5 | Rank 6 | Rank 7 | Rank 8 | Rank 9 | Rank 10 |
|---|---|---|---|---|---|---|---|---|---|---|
| A | 0.001 | 0.001 | 0.001 | 0.001 | 0.001 | 0.001 | 0.001 | 0.001 | 0.001 | 0.001 |
| C | 0.208 | 0.001 | 0.001 | 0.001 | 0.001 | 0.001 | 0.001 | 0.001 | 0.001 | 0.001 |
| D | 0.783 | 0.001 | 0.001 | 0.001 | 0.001 | 0.001 | 0.001 | 0.001 | 0.001 | 0.001 |

| Subject | Rank 11 | Rank 12 | Rank 13 | Rank 14 | Rank 15 | Rank 16 | Rank 17 | Rank 18 | Rank 19 | Rank 20 |
|---|---|---|---|---|---|---|---|---|---|---|
| A | 0.001 | 0.001 | 0.001 | 0.001 | 0.001 | 0.001 | 0.001 | 0.001 | 0.001 | 0.001 |
| C | 0.001 | 0.001 | 0.001 | 0.001 | 0.001 | 0.001 | 0.001 | 0.001 | 0.001 | 0.001 |
| D | 0.001 | 0.001 | 0.001 | 0.001 | 0.001 | 0.001 | 0.001 | 0.001 | 0.001 | 0.001 |

(b) $CC_{norm}$ differences between low-rank models vs full linear rank regression are significant at 0.001 for all models except for Subject C and D rank-1 models. Shown above are the estimated p-values from a bootstrapped test performance measure. To estimate the p-value, test data was split into sequential blocks of 10 seconds, sampled from with replacement and used to construct a new test dataset. The performance difference between the low-rank model and full rank model were calculated for each new test set. The p-value is then the ratio of performances less than 0 to total bootstraps, in this case 1000

Figure 11: Component scaling over factors and significance difference tests between rank models and full rank ridge regression.

| Subject | Rank 1 | Rank 2 | Rank 3 | Rank 4 | Rank 5 | Rank 6 | Rank 7 | Rank 8 | Rank 9 | Rank 10 |
|---|---|---|---|---|---|---|---|---|---|---|
| A | 0.001 | 0.001 | 0.001 | 0.001 | 0.001 | 0.001 | 0.001 | 0.001 | 0.001 | 0.001 |
| C | 0.001 | 0.001 | 0.001 | 0.001 | 0.001 | 0.001 | 0.001 | 0.001 | 0.001 | 0.001 |
| D | 0.001 | 0.001 | 0.001 | 0.001 | 0.001 | 0.001 | 0.001 | 0.001 | 0.001 | 0.001 |

| Subject | Rank 11 | Rank 12 | Rank 13 | Rank 14 | Rank 15 | Rank 16 | Rank 17 | Rank 18 | Rank 19 | Rank 20 |
|---|---|---|---|---|---|---|---|---|---|---|
| A | 0.001 | 0.001 | 0.001 | 0.001 | 0.001 | 0.001 | 0.001 | 0.001 | 0.001 | 0.001 |
| C | 0.001 | 0.001 | 0.001 | 0.001 | 0.001 | 0.001 | 0.001 | 0.001 | 0.001 | 0.001 |
| D | 0.001 | 0.001 | 0.001 | 0.001 | 0.001 | 0.001 | 0.001 | 0.001 | 0.001 | 0.001 |

Figure 12: $CC_{norm}$ performance is above chance for all models at 0.001. Shown above are the estimated p-values from a permuted test performance measure. To estimate p-values, we first split our test MEG data into sequential blocks of 10 seconds. Then for each permutation sample, we shuffled the blocks to create a new test set and computed the performance of the model on it. Using all permutations (in this case 1000), we construct the null distribution that our model is equal to a chance model. Then, we obtain the p-value as the probability that the null distribution is greater than the test performance of the model.

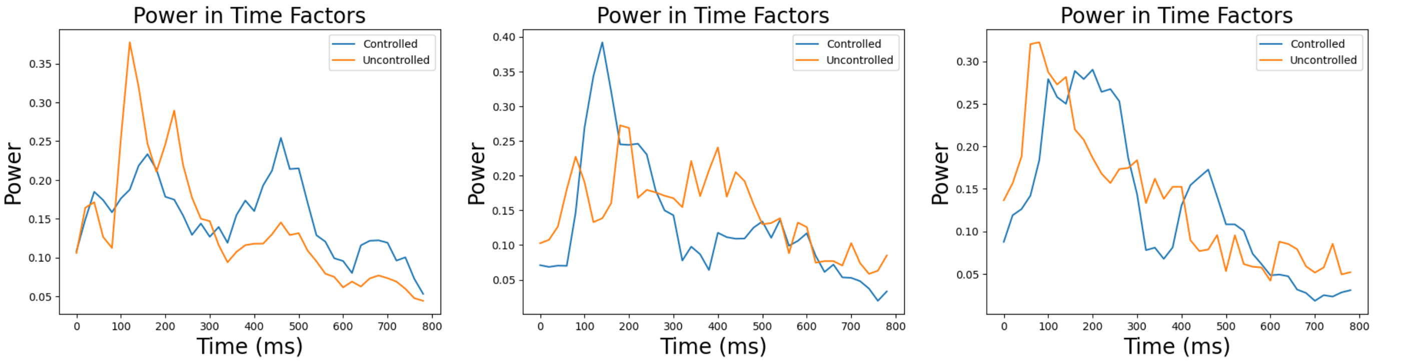


Figure 13: The power in the time factors shifts to be later in subjects A and D after subtracting controls. In subjects A and D, the power is approximately monotonic before subtracting controls, peaking at around 150ms. After control subtraction, double peaks form, one early peak at around 150-200ms followed by a later peak around 400-500ms. Subject C exhibits this double peak behavior, however after controls, it becomes more peaked around 150ms. *Left*: Subject A. *Middle*: Subject C. *Right*: Subject D.

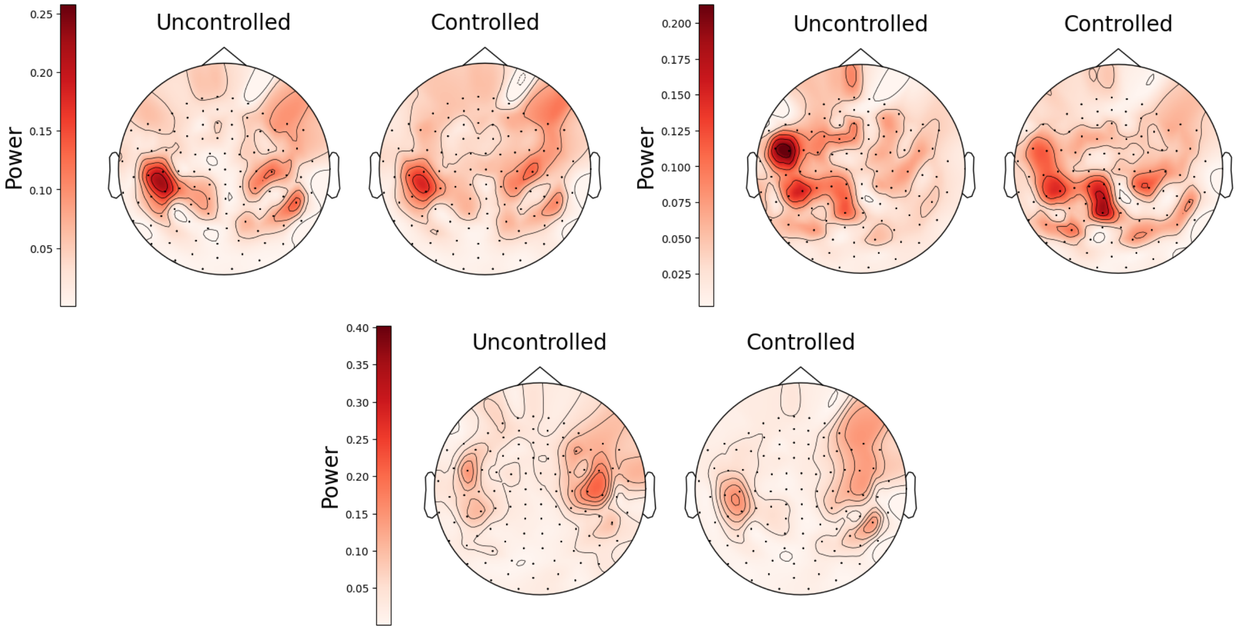


Figure 14: Power in channel factors before and after subtracting controls. Generally, across subjects, channel factors shift to become less concentrated over auditory areas, affecting channels more globally following control subtraction. *Top left*: Subject A. *Top right*: Subject C. *Bottom middle*: Subject D.
